# Masking phosphatidylserine prevents neuronal loss in two distinct *Drosophila* models of neurodegeneration

**DOI:** 10.64898/2026.07.01.735827

**Authors:** Naden Khateb, Miri Shwartsburd, Eden Grig, Shelly Vogelesang-Ganon, Tsneem Fauzi, Malak Ayoub, Ketty Hakim-Mishnaevski, Estee Kurant

**Affiliations:** The Herta & Paul Amir School of Medicine and the Department of Human Biology, Faculty of Natural Sciences, University of Haifa, Haifa 34988, Israel

**Keywords:** CNS, *Drosophila*, glia, Lactadherin, MFG-E8, phagocytosis, phagoptosis, SkpA, Huntington’s disease, neurodegeneration

## Abstract

Neuronal loss is a hallmark of neurodegenerative diseases. Phosphatidylserine (PS), a key ‘eat me’ signal, is exposed on stressed viable neurons, triggering their premature phagocytosis by activated glia. We investigated whether PS masking could serve as a universal strategy to prevent neuronal loss in two distinct *Drosophila* models of neurodegeneration: an adult-stage-specific knockdown of *skpA* and a Huntington’s disease model initiated during embryogenesis. Both models exhibit neuronal loss, motor dysfunction, and reduced lifespan. To mask PS, we used a truncated form of MFG-E8, a glycoprotein that binds PS without promoting engulfment. PS masking preserved two neuronal populations in both models, indicating that these neurons were eliminated alive via phagoptosis. Motor function and lifespan were improved to varying degrees, depending on the timing and severity of neuronal damage. These findings reveal that aberrant glial phagocytosis contributes to neuronal vulnerability and identify PS masking as a promising therapeutic approach for neurodegenerative diseases.

**Significance Statement:** Neuronal loss is a defining feature of neurodegenerative diseases, yet its underlying mechanisms remain incompletely understood. Here, we demonstrate in two *Drosophila* models of neurodegeneration that stressed but viable neurons are prematurely eliminated by glial phagocytosis through phosphatidylserine (PS) exposure. By masking PS with a truncated form of MFG-E8, we prevented neuronal loss, improved motor performance, and extended lifespan, highlighting PS-dependent removal of live neurons as a critical contributor to neurodegeneration. Our findings provide the first *in vivo* evidence that PS masking protects neurons in distinct neurodegenerative contexts, offering a broadly applicable strategy for therapeutic intervention. This work positions aberrant glial phagocytosis as a disease-driving mechanism and establishes *Drosophila* as a powerful model for dissecting neuron-glia interactions in neurodegeneration.

## Introduction

Neurodegenerative diseases are characterized by progressive neuronal loss in the central nervous system (CNS), leading to cognitive decline, motor dysfunction, and premature death. Despite their increasing prevalence, no curative therapies are currently available (1). Recent studies have highlighted the pivotal role of phagocytic glia in neurodegeneration (2, 3). These glial cells contribute to neuronal loss by engulfing stressed viable neurons through a phosphatidylserine (PS)-dependent mechanism known as ‘phagoptosis’ (4–6). Consequently, masking PS has emerged as a promising strategy to rescue stressed neurons and delay the progression of neurodegeneration (6). Yet, the potential of PS masking in preventing neurodegeneration has never been tested *in vivo*. Moreover, the selective vulnerability of neuronal populations, often reflected by specific neuronal loss in different disorders, raises the important question of how selective and context-dependent glial phagocytosis is, and whether it contributes differently to neuronal loss and disease progression.

Model organisms, particularly the fruit fly *Drosophila melanogaster*, provide an invaluable platform for investigating the cellular and molecular mechanisms underlying neurodegenerative pathologies due to their relatively simple brain structure, accessibility to imaging, short lifespan, and advanced genetic tools (7–10). In *Drosophila*, phagocytic glia efficiently clear apoptotic neurons during development, a process mediated by three main receptors: Santa-maria, SIMU, and Draper (11–13). Both SIMU and Draper have been demonstrated to bind PS (14, 15). Notably, overexpression of these receptors in adult brain glia is sufficient to induce PS-dependent neuronal loss, accompanied by motor deficits and reduced lifespan (16).

In this study, we used two *Drosophila* models of neurodegeneration to investigate the impact of PS masking on neuronal loss. The first model is based on pan-neural knockdown of *skpA* (a *Drosophila* homolog of human *skp1*), a component of the ubiquitin E3 ligases (17, 18), specifically in the adult brain, leading to accumulation of ubiquitinated protein aggregates, neuronal loss, motor dysfunction, and reduced lifespan (19, 20). Human Skp1, which shares 76% protein homology with *Drosophila* SkpA has been shown to be decreased in patients with Parkinson’s disease (PD) (21, 22), and its silencing was used to model PD in cell lines and mice (23, 24). The second model involves neuronal expression of the human mutant Huntingtin (Htt) protein containing an expanded polyglutamine (polyQ) tract of 128 repeats (128Q). A polyQ length exceeding 35 repeats is known to cause autosomal dominant Huntington’s disease (HD) (25). Various *Drosophila* models of HD have been developed and used in previous studies (26). In our model, Htt expression begins during embryogenesis and leads to progressive neuronal loss, motor dysfunction, and reduced lifespan, recapitulating key features of HD pathogenesis.

To mask PS in both models, we used Milk Fat Globule-Epidermal Growth Factor 8 (MFG-E8), also known as Lactadherin (Lact), a mammalian glycoprotein secreted by activated macrophages or microglia (27). MFG-E8 acts as a bridging molecule, binding PS on apoptotic cells and integrins on phagocyte membranes, thereby enhancing the engulfment of apoptotic debris or viable cells in inflammation (28, 29). The truncated form of human MFG-E8 (LactC1C2), which lacks the integrin-binding region (30) was used to mask PS to prevent glial phagocytosis.

Our data demonstrate that neuronal loss in the adult brain caused by *skpA* RNAi knockdown or by expanded human Htt expression can be fully prevented by the truncated LactC1C2. In contrast, LactC1C2 did not rescue neuronal loss during development of HD flies, suggesting different mechanisms of Htt-induced effect at this stage. Importantly, motor ability and survival rate depended not only on neuronal persistence but also on proper neuronal function. We found that, in the *skpA* RNAi model, LactC1C2-rescued neurons did not regain full functionality, whereas in the HD model, both motor ability and lifespan were restored close to control levels. These findings from two distinct models reveal that in the diseased adult brain stressed neurons are removed by phagocytic glia, and can be protected by masking PS.

## Results

### Generation of a *skpA*-based model of neurodegeneration in adult *Drosophila*

We previously identified the neuroprotective role of the *Drosophila skpA* gene and demonstrated that pan-neural knockdown of *skpA* by RNAi in the adult fly brain leads to the loss of dopaminergic (DA) neurons, accompanied by reduced climbing ability and decreased survival rates (19). Based on these findings obtained with the *elavGal4* pan-neural driver, we established an improved fly model of neurodegeneration by employing the stronger pan-neural *nsybGal4* driver to express *skpA* RNAi (31–33) only at the adult stage. For this, we utilized a temperature-sensitive TubGal80 repressor of Gal4, which is active at temperatures lower than 23°C and inactive at 29°C, allowing temporal regulation of Gal4-induced expression (*Gal4/Gal80ts* system) (34). Newly eclosed flies carrying *nsybGal4*-driven *skpA* RNAi and *tubGal80* were immediately transferred from 20°C to 29°C. This shift deactivated Gal80 repression, enabling Gal4 activity and subsequent reduction of SkpA levels (Fig. 1A).

**Figure 1.**
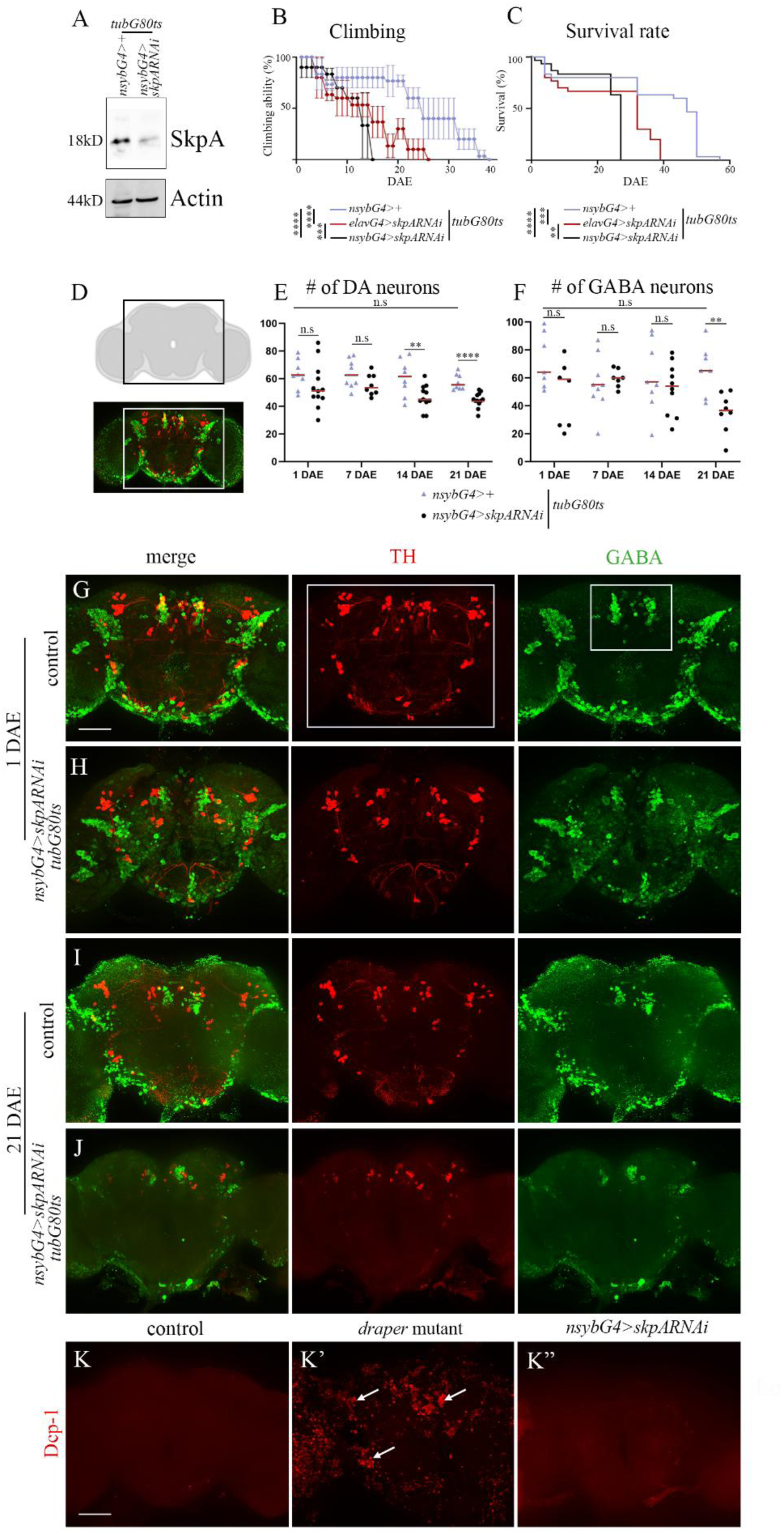
*skpA* RNAi knockdown in adult brain neurons reduces motor ability and lifespan accompanied by loss of DA and GABA neurons. (A) Western blot analysis of head lysates from wild type and *skpA* RNAi (*tubGal80ts/+;nsybGal4/UASskpARNAi*) flies at 21 DAE with anti-SkpA and anti-Actin antibodies. (B, C) Climbing ability (B) and survival rates (C) of flies expressing *skpA* RNAi driven by *elavGal4* (*elavGal4/+;tubGal80ts/UASskpARNAi*) (red) or by *nsybGal4* (*tubGal80ts/+;nsybGal4/UASskpARNAi*) (black) compared with control flies (*nsybGal4/+*) (purple) at 29℃ from 1 DAE. Data are represented as mean ± SEM, n (number of vials, starting with 10 female flies in each) = 3. ****p < 0.0001, ***p < 0.001, **p < 0.01. (D) Diagram and immunofluorescent image of the brain immunostained with anti-TH (DA neurons, red) and anti-GABA (GABA neurons, green) antibodies. The neurons located in the designated area of the brain (black and white frames). (E, F) Mean total number of DA neurons (E) and GABA neurons (F) in dissected brains of flies at different age (1,7,14,21 DAE) ± SEM, n (number of brains) ≥7. ****p < 0.0001, **p < 0.01, n.s = non-significant. (G-J) Representative images of projections of Z-stacks of the posterior part (∼45 µm) of whole-mount brains from females maintained at 29℃. DA neurons labeled with anti-TH (red), GABA neurons labeled with anti-GABA (green). Bar, 50 µm. (G) control (*tubGal80ts/+; nsybGal4/+*) 1 DAE. (H) *skpA* RNAi knockdown (*tubGal80ts/+;nsybGal4 UASskpARNAi*) 1 DAE. (I) control (*tubGal80ts/+; nsybGal4/+*) 21 DAE. (J) *skpA* RNAi knockdown (*tubGal80ts/+;nsybGal4/UASskpARNAi*) 21 DAE. (K-K”) Immunostaining with anti-Dcp-1 (red) of 21 DAE brains. (K) control (*tubGal80ts/+;nsybGal4/+*), (K’) *draper* mutant showing accumulation of apoptotic debris (arrows), (K”) *skpA* RNAi knockdown (*tubGal80ts/+;nsybGal4/UASskpARNAi*), no detectible Dcp-1 staining. Bar, 50 µm.

### RNAi-mediated *skpA* knockdown in adult brain neurons causes neuronal loss

To evaluate neurodegeneration in flies, three main assays are commonly applied: 1) assessing motor ability through climbing behavior (negative geotaxis), 2) measuring lifespan, and 3) labeling and quantifying neurons in the brain (16, 19). Transgenic flies expressing *skpA* RNAi under *nsybGal4* only in adulthood exhibited significantly greater locomotor deficits and shorter lifespans compared to flies expressing *skpA* RNAi with *elavGal4* and control flies outcrossed to *w1118* (Fig. 1B, C), suggesting strong neurodegenerative defects.

To monitor neuronal loss in adult fly brains, we focused on two distinct neuronal populations in the central brain area (Fig. 1D): DA neurons (Fig. 1E) and GABAergic (GABA) neurons (Fig. 1F). These neuronal populations were labeled with anti-Tyrosine hydroxylase (TH) and anti-GABA antibodies, respectively (Fig. 1G-J). Neuronal counts were conducted at four time points: 1, 7, 14 and 21 days after eclosion (DAE) in specific designated regions of the central brain (Fig. 1G) using Imaris (Oxford Instruments) software (Fig. 1E, F). Compared to control brains, a progressive decrease in the number of DA and GABA neurons was observed in brains with reduced SkpA expression (Fig. 1E, F). At 14 DAE significant reduction in DA neuronal number was observed with further decline at 21 DAE (Fig. 1E). GABA neuronal number stayed similar to control at 14 DAE but was significantly decreased at 21 DAE (Fig. 1F) suggesting differential sensitivity of DA and GABA neurons to the SkpA reduced levels. These findings highlight the neuroprotective role of SkpA and validate the adult-specific *skpA* RNAi neuronal knockdown as a suitable model for studying the mechanisms underlying loss of different types of neurons in the adult brain.

### Neuronal loss in *skpA* knockdown flies is not mediated by apoptosis

To determine whether neuronal loss in *skpA* knockdown flies is caused by apoptosis, we examined caspase activation using immunostaining with an antibody against cleaved *Drosophila* Caspase-1 (Dcp-1). As expected, control adult brains showed no detectable signal, consistent with the low levels of apoptosis reported in the adult *Drosophila* brain (35) (Fig. 1K, S1). In contrast, *draper* mutant brains, used as a positive control, exhibited strong Dcp-1 staining, reflecting the accumulation of uncleared apoptotic neurons (Fig. 1K’, S1). Notably, brains from *skpA* knockdown flies also lacked the Dcp-1 signal (Fig. 1K”, S1), resembling the pattern seen in controls. These findings suggest that the observed neuronal loss in *skpA* knockdown flies is not mediated by apoptosis.

### PS exposure and Draper expression are increased in the *skpA* RNAi-induced model

In mammals, activated microglia have been shown to phagocytose stressed viable neurons exposing PS on their surfaces during inflammation (4, 5, 36). This process, called phagoptosis, points to an active role of phagocytosis in the cell death process. In this line, we previously demonstrated a causative role of glial phagocytosis in neuronal loss in the *Drosophila* adult brain, which was partially prevented by the secreted human MFG-E8 (16). To visualize and assess PS levels in the *skpA* RNAi model, we stained dissected adult brains with a fluorescent Annexin V, which binds PS. We performed this staining in control brains and brains of *skpA* RNAi-induced neurodegeneration model at 1 and 21 DAE (Fig. 2A-C). Low levels of Annexin V labeling were found in control brains at 1 and 21 DAE (Fig. 2A-C), as well as in 1 DAE *skpA* RNAi flies (Fig. 2A’, C). In contrast, *skpA* RNAi brains at 21 DAE exhibited higher levels of Annexin V labeling (Fig. 2B’, C), consistent with increased PS exposure. In addition, to assess phagocytic activity of glia, we measured Draper levels in the *skpA* RNAi model flies and compared this to control (Fig. 2D-F). It was previously shown that under normal conditions, Draper declines with age (37). Consistent with this, our data show that Draper levels decrease at 21 DAE compared to 1 DAE in control flies (Fig. 2D, E, F). However, in the *skpA* model flies Draper levels are much higher at 21 DAE compared to 1 DAE and control flies (Fig. 2D’, E’, F). These findings suggest that live neurons stressed by *skpA* depletion expose more PS, which is recognized by Draper expressed in higher levels in glia leading to their phagocytosis.

**Figure 2.**
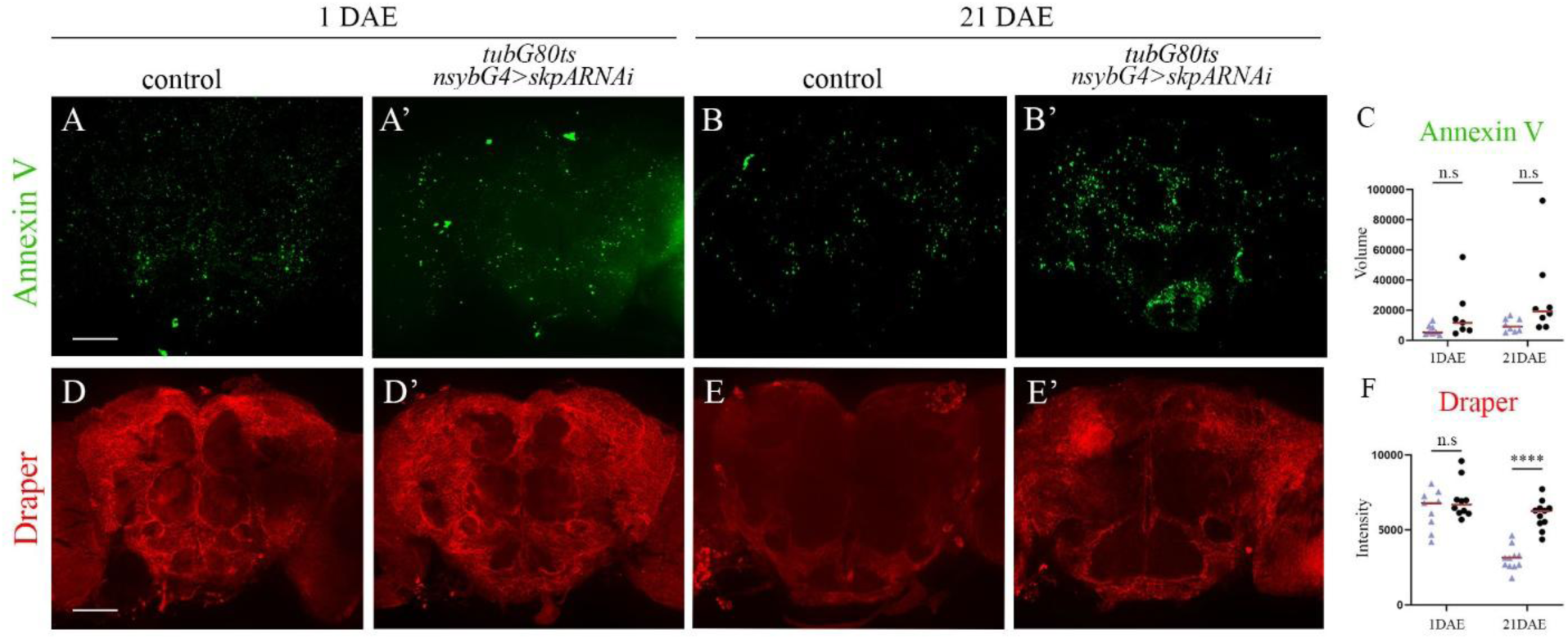
Increased PS exposure and Draper expression in the *skpA* RNAi-induced model. (A–B’, D–E’) Representative projections of Z-stacks of the posterior region (∼45 µm) of whole-mount brains from female flies maintained at 29°C. (A–B’) PS is visualized using fluorescent Annexin V (green). (D–E’) Draper expression is detected by immunostaining with anti-Draper (red). (A, B, D, E) Control (*tubGal80ts/+; nsybGal4/+*). (A′, B′, D′, E′) *skpA* RNAi knockdown (*tubGal80ts/+;nsybGal4/UASskpARNAi*). Scale bar, 50 µm. (C) Mean total volume of Annexin V–positive signal within whole brain± SEM; n (number of brains) ≥ 7. n.s = not significant. (F) Mean intensity of anti-Draper staining within whole brain ± SEM; n (number of brains) ≥ 9. ****p < 0.0001, n.s = not significant).

### GFP-tagged LactC1C2 is secreted from adult brain neurons

Based on these findings, we hypothesized that if stressed neurons are phagocytosed while still viable, masking PS could rescue neuronal loss in the *skpA* RNAi-induced neurodegeneration model by inhibiting phagocytosis. To test this, we utilized a GFP-tagged truncated version of MFG-E8 (LactC1C2-GFP), which binds PS but not integrins (30), preventing engulfment (Fig. 3A). Since LactC1C2-GFP is a secreted molecule, we used the pan-neural *nsybGal4* driver to express and distribute *lactC1C2-GFP* throughout the brain starting from 1 DAE employing the Gal4/Gal80ts system. As shown in Figures 3B and 3C, LactC1C2-GFP was detected in the adult brains of these flies. We also examined LactC1C2-GFP expression in the *skpA* RNAi background (Fig. S2) and detected GFP in patches on neuronal surfaces suggesting that secreted LactC1C2-GFP binds PS exposed on neurons (Fig. S2).

**Figure 3.**
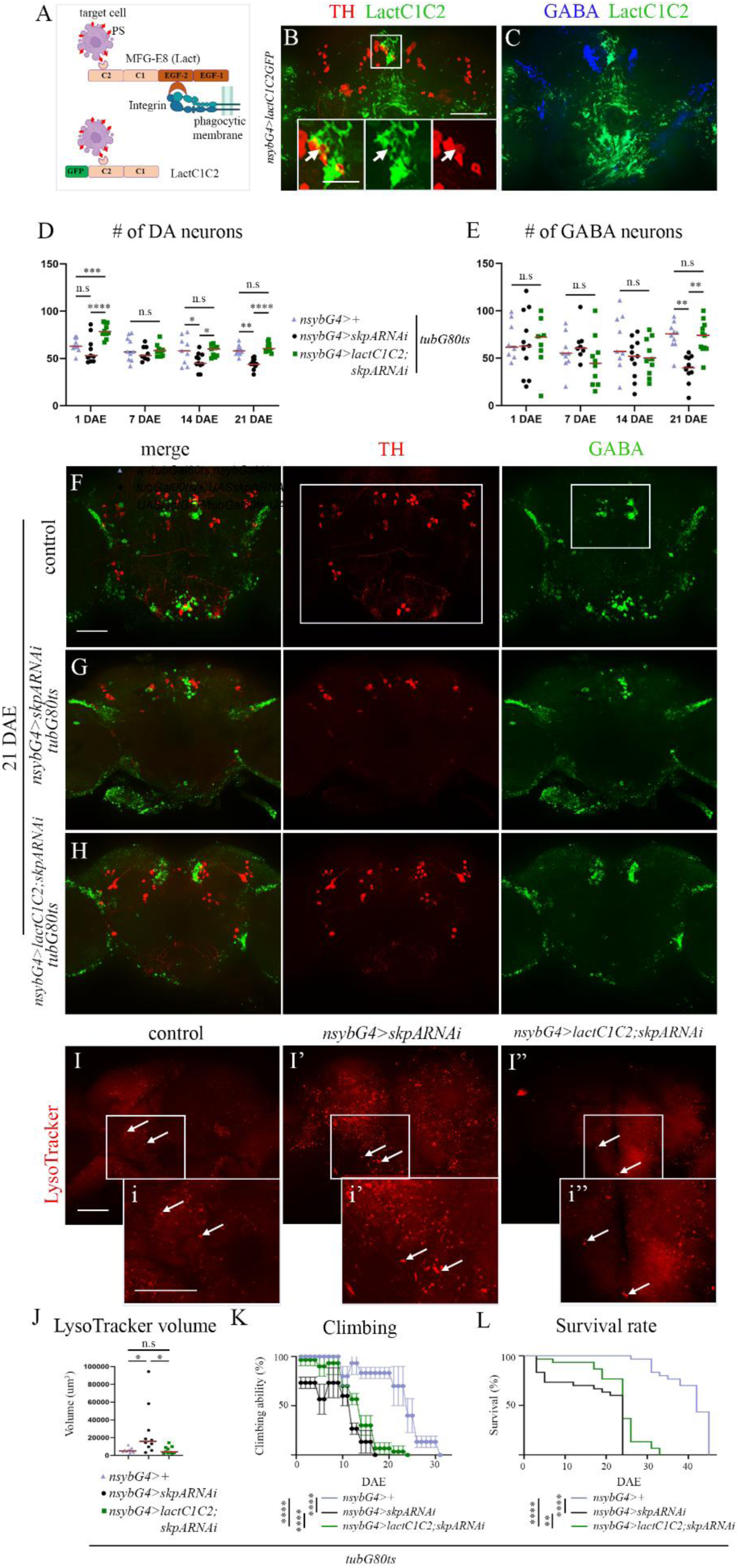
LactC1C2-GFP secreted from neurons rescues DA and GABA neurons and partially rescues motor ability and extends lifespan of *skpA* knockdown flies. (A) Schematic representation of 4 different domains in the Lact protein, depicting regions responsible for binding to PS on target cell and for interaction with integrins on phagocytic membrane. GFP-tagged truncated form of Lact (LactC1C2-GFP). (B, C) Projections of Z-stacks of the posterior part (∼45 µm) of whole-mount female brain carrying LactC1C2-GFP (*nsybG4>lactC1C2-GFP)* maintained at 29℃ 10 DAE. DA neurons labeled with anti-TH (red), GABA neurons labeled with anti-GABA (blue), LactC1C2-GFP (green). Insets in B enlarge the rectangle area in separate channels, arrows depict TH-labeled neurons with GFP signal accumulating on their surfaces. Bar, 20 µm. (D, E) Mean total number of DA neurons (D) and GABA neurons (E) in dissected brains of flies at different age (1,7,14,21 DAE) ± SEM, n (number of brains) ≥8. ****p < 0.0001, ***p<0.001, **p < 0.01, *p<0.05, n.s = non-significant. (F-H) Projections of Z-stacks of the posterior part (∼45 µm) of whole-mount 21 DAE brains maintained at 29℃. DA neurons labeled with anti-TH (red), GABA neurons labeled with anti-GABA (green). Bar, 50 µm. (F) control (*tubGal80ts/+; nsybGal4/+*). (G) *skpA* RNAi knockdown (*tubGal80ts/+;nsybGal4/ UASskpARNAi*). (H) *lactC1C2-GFP* and *skpA* RNAi (*tubGal80ts/UASlactC1C2-GFP;nsybGal4/ UASskpARNAi*). (I-I”) LysoTracker staining (red) of 21 DAE brains. (I) control (*tubGal80ts/+;nsybGal4/+*), (I’) *skpA* RNAi knockdown (*tubGal80ts/+;nsybGal4/UASskpARNAi*), (I”) *lactC1C2-GFP* and *skpA* RNAi (*tubGal80ts/UASlactC1C2-GFP;nsybGal4/UASskpARNAi*). (i-i”) close-ups of the rectangular regions shown in I–I’’. Arrows indicate LysoTracker-positive phagosomes. Bar, 50 µm. (J) Mean total volume of LysoTracker-positive area within whole brain ± SEM, n (number of brains) = 10. *p<0.05, n.s = non-significant. (K, L) Climbing ability (K) and survival rates (L) of flies expressing *skpA* RNAi (*tubGal80ts/+;nsybGal4/UASskpARNAi*) (black), *lactC1C2-GFP* and *skpA* RNAi (*tubGal80ts/UASlactC1C2-GFP;nsybGal4/UASskpARNAi*) (green) and control flies (*tubGal80ts/+;nsybGal4/+*) (purple). Data are represented as mean ± SEM, n (number of vials, starting with 10 female flies in each) = 3. ****p < 0.0001, **p < 0.01.

### PS masking rescues DA and GABA neurons in the *skpA* RNAi-induced model

To evaluate the impact of PS masking by LactC1C2-GFP on DA and GABA, we quantified their numbers (Fig. 3D, E) using anti-TH and anti-GABA antibodies to label DA and GABA neurons, respectively (Fig. 3F-H). These markers label live neurons but not dead cells. Neuronal numbers were counted at 1, 7, 14 and 21 DAE in specific designated regions of the central brain (Fig. 3F) using Imaris (Oxford Instruments) software (Fig. 3D, E). Remarkably, neuronal expression of LactC1C2-GFP in the adult brain completely rescued the loss of DA and GABA neurons induced by *skpA* RNAi compared to flies expressing only *skpA* RNAi (Fig. 3D, E). For GABA neurons, the average number in the designated area was significantly higher in flies expressing LactC1C2-GFP (mean = 62.5) than in flies expressing *skpA* RNAi alone (mean = 43.5) at 21 DAE and was not significantly different from control flies (mean = 69.4) (Fig. 3E). Similarly, for DA neurons, the average number in flies expressing LactC1C2-GFP in adult brain neurons at 1 DAE (mean = 77.8) was significantly higher than in flies expressing *skpA* RNAi alone (mean = 56) and even higher than in control flies (mean = 64.9) (Fig. 3D). However, no significant difference was found between control and LactC1C2-GFP expressing *skpA* knockdown flies at 21 DAE (Fig. 3D), indicating that PS masking effectively prevented the removal of neurons. Collectively, these results are consistent with the model that PS exposure on neuronal surfaces triggers glial phagocytosis, potentially contributing to the loss of stressed neurons and exacerbating neurodegeneration, a process that can be prevented by masking PS by LactC1C2-GFP.

### Enhanced phagocytosis in *skpA* knockdown fly brains is reduced by LactC1C2

To assess phagocytic activity in the adult *Drosophila* brain, we labeled lysosomes and phagolysosomes using LysoTracker, a red fluorescent marker of acidic compartments, in flies at 21 DAE. In dissected brains of control flies, we observed small LysoTracker-positive vesicles, consistent with basal lysosomal activity (Fig. 3I, J). In contrast, brains from *skpA* RNAi flies exhibited a markedly increased number of enlarged LysoTracker-positive vesicles (Fig. 3I’, J). To determine which cells are phagocytic and contain large LT-positive vesicles, we selectively labeled glial or neuronal membranes (Fig. S3). We found that all large vesicles were located within glial cells labeled with an anti-Draper antibody (Fig. S3A-C”). In contrast, when neuronal membranes were labeled using *nsybGal4>CD8GFP*, only small LT-positive vesicles were observed within neurons (Fig. S3D-F”). The increased number of LysoTracker-positive vesicles is consistent with enhanced glial phagocytic activity. Notably, in *skpA* RNAi flies expressing LactC1C2-GFP, LysoTracker staining revealed fewer and smaller vesicles, resembling the pattern observed in controls (Fig. 3I”, J). These findings suggest that the enhanced phagocytic activity in *skpA* knockdown brains is attenuated by PS masking via LactC1C2-GFP expression, likely by preventing phagoptosis of live stressed neurons that expose PS on their surface.

### PS masking partially improves motor ability and survival rate in *skpA* knockdown flies

To assess whether the rescued neurons in *skpA* knockdown flies expressing LactC1C2-GFP remain functional, we evaluated the effect of PS masking on flies’ motor ability and lifespan. In the climbing assay, flies expressing LactC1C2-GFP in adult brain neurons exhibited improved motor performance compared to those with *skpA* RNAi alone, although they still displayed locomotor deficits relative to control flies (Fig. 3K). Notably, *skpA* knockdown flies showed reduced climbing activity from 1 DAE, whereas those expressing LactC1C2-GFP maintained performance until 10 DAE before experiencing decline (Fig. 3K). Similarly, lifespan analysis revealed that LactC1C2-GFP expression extended survival of *skpA* knockdown flies, although their lifespan remained shorter than that of control flies (Fig. 3L). *skpA* knockdown flies died after 24 days, whereas those co-expressing LactC1C2-GFP survived up to 33 days, and control flies lived up to 45 days (Fig. 3L). These findings demonstrate that PS masking with LactC1C2-GFP improves motor function and extends lifespan in *skpA* knockdown flies, albeit not at the wild type level.

### Neuronal expression of human Htt128Q reduces motor activity and shortens lifespan

To compare the effect of masking PS on neuronal loss between different modes of neurodegeneration, we employed the HD *Drosophila* model of neurodegeneration (26), which differs from the *skpA* RNAi-induced model in two key aspects. First, it involves pan-neural expression of human Htt128Q (*UAShtt128Q*) using the *nsybGal4* driver. Second, Htt128Q expression begins at embryogenesis, mimicking the human condition in which the expanded mutant Htt is present throughout life. To assess adult brain function in HD model flies, we evaluated their climbing ability and survival rate (Fig. 4A, B). HD flies showed a marked motor deficit, losing climbing ability within two weeks after eclosion, in contrast to control flies that retained high climbing ability for over 50 days at 25°C (Fig. 4A). Lifespan was also significantly reduced in HD flies, with the majority dying within three weeks post-eclosion, compared to control flies which survived for over two months (Fig. 4B). These findings demonstrate that neuronal expression of Htt128Q using the *nsybGal4* driver results in progressive motor dysfunction and premature death, recapitulating key features of neurodegenerative disease in *Drosophila*.

**Figure 4.**
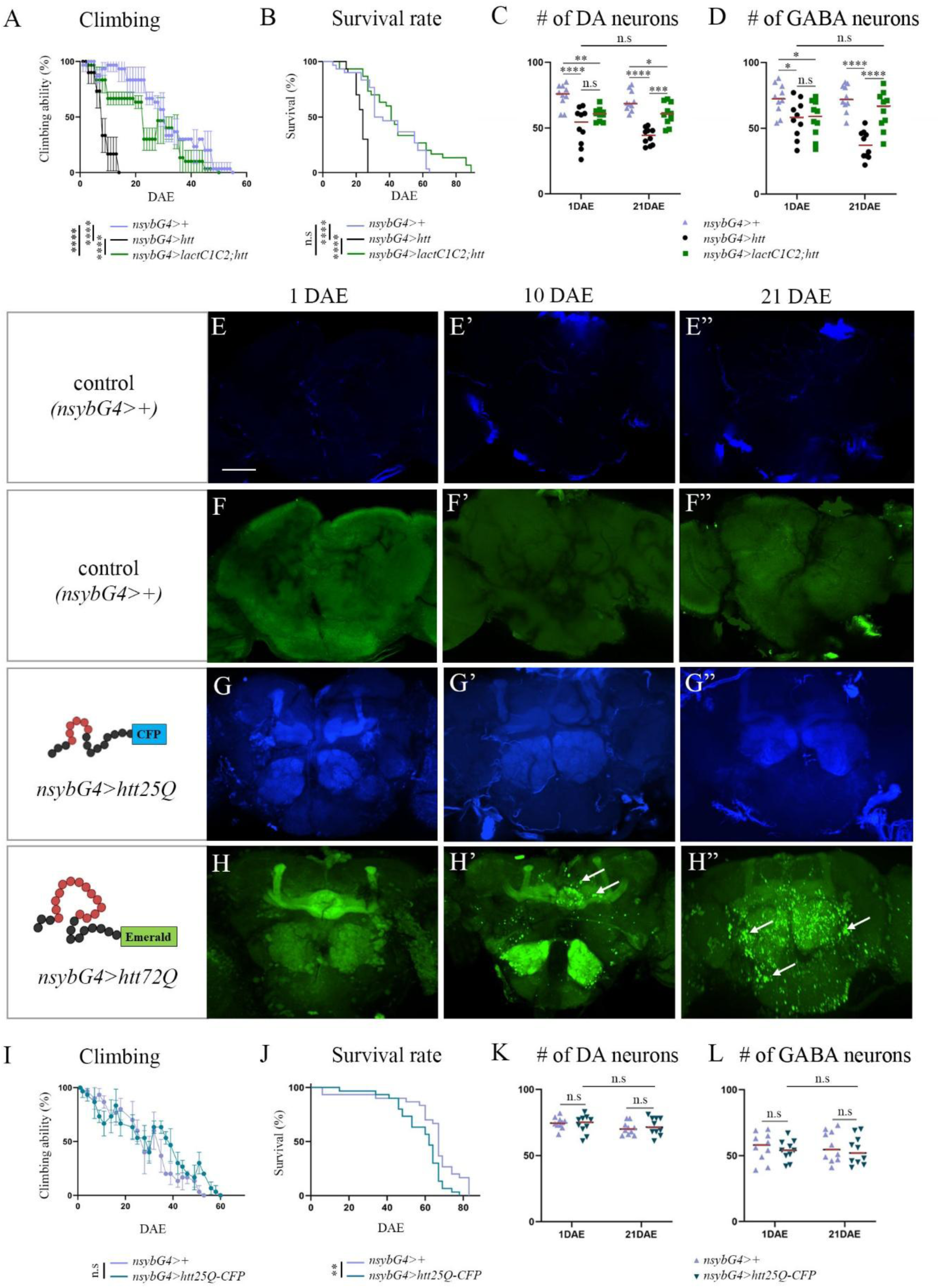
Pan-neural expression of human expanded Htt reduces motor ability and lifespan accompanied by loss of DA and GABA neurons, which is prevented by LactC1C2-GFP. (A, B) Climbing ability (A) and survival rates (B) of flies expressing human Htt128Q (*nsybGal4/UAShtt*) (black), human Htt128Q and LactC1C2-GFP (*UASlactC1C2-GFP/+;nsybGal4/UAShtt*) (green), and control flies (*nsybGal4/+*) (purple). Data are represented as mean ± SEM, n (number of vials, starting with 10 female flies in each) = 3. ****p < 0.0001, n.s = non-significant. (C, D) Mean total number of DA (C) and GABA neurons (D) in dissected brains of flies expressing human Htt128Q (*nsybGal4/UAShtt*) (black), human Htt128Q and LactC1C2-GFP (*UASlactC1C2-GFP/+;nsybGal4/UAShtt*) (green), and control flies (*nsybGal4/+*) (purple) at 1 DAE and 21 DAE ± SEM, n (number of brains) = 10, ****p < 0.0001, ***p < 0.001, **p < 0.01, *p < 0.05, n.s = non-significant. (E-H”) Microscope images of the posterior part (∼45 µm) of whole-mount female brains of (E-F”) control (*nsybGal4/+*), (G-G”) *UAShtt25Q-CFP/+;nsybGal4/+*, and (H-H”) *UAShtt72Q-Emerald/+;nsybGal4/+* at 1, 10 and 21 DAE. (H’, H”) Arrows depict aggregates of Htt72Q-Emerald. Bar, 50 µm. (I, J) Climbing ability (I) and survival rates (J) of flies expressing Htt25Q-CFP *(UAShtt25Q-CFP+;nsybGal4/+*) (green) and control flies (*nsybGal4/+*) (purple). Data are represented as mean ± SEM, n (number of vials, starting with 10 female flies in each) = 3. **p < 0.01, n.s = non-significant. (K, L) Mean total number of DA (K) and GABA neurons (L) in dissected brains of flies expressing Htt25Q-CFP *(UAShtt25Q-CFP/+;nsybGal4/+*) (green) and control flies (*nsybGal4/+*) (purple) at 1 DAE and 21 DAE ± SEM, n (number of brains) = 10, n.s = non-significant.

### The number of DA and GABA neurons is significantly reduced in HD model flies

To evaluate neuronal loss in the brains of adult HD flies, we quantified DA and GABA neuronal populations in the central brain, labeled with anti-TH and GABA respectively, as in the *skpA* RNAi-induced model. Brains were dissected, immunostained, and imaged at 1 DAE and 21 DAE, just before the flies died. Both neuronal populations were affected by the expression of human expanded Htt, though to differing extents. At 1 DAE, the number of DA neurons in HD flies was slightly but significantly reduced compared to that of control flies (Fig. 4C), whereas the number of GABA neurons was already much strongly reduced at 1 DAE compared to controls (Fig. 4D). These findings indicate that some neuronal loss occurred during development, with DA and GABA neurons exhibiting differential vulnerability to aberrant Htt protein expression. By 21 DAE, both DA and GABA neuronal populations showed a further decreased number in HD flies compared to controls (Fig. 4C, D) highlighting the pronounced and damaging effect of the expanded human Htt protein on these two neuronal cell types in the adult brain.

### A fluorescently-tagged expanded human Htt forms aggregates in the adult fly brain

In humans, the tendency of mutant Htt protein to aggregate and form inclusions is a key hallmark of HD (25). To confirm that expanded Htt forms aggregates, we examined the expression and distribution of human Htt protein in the adult *Drosophila* brain using two transgenic lines: one encoding Cerulean Fluorescent Protein (CFP)-tagged non-expanded 25Q Htt (*UAShtt25Q-CFP*) and the other encoding Emerald-tagged expanded 72Q Htt (*UAShtt72Q-Emerald*) (38). These constructs were expressed pan-neuronally from embryogenesis using the *nsybGal4* driver. Adult brains were dissected and analyzed at 1, 10, and 21 DAE (Fig. 4E-H”). In control brains, only weak autofluorescent background signals were observed in both fluorescence channels across all time points (Fig. 4E-F”). In brains expressing the shorter Htt25Q-CFP, fluorescence was homogeneously distributed without visible aggregates at all time points (Fig. 4G-G”), indicating normal Htt distribution. To confirm that neuronal expression of the short Htt does not induce neurodegeneration, we performed all three assays in flies expressing Htt25Q-CFP starting during embryogenesis. Our data demonstrate that neuronal expression of the short form of human Htt does not affect neuronal numbers and has no impact on climbing performance or survival (Fig. 4I-L).

In contrast, brains expressing Htt72Q-Emerald showed a homogeneous distribution of Emerald at 1 DAE (Fig. 4H), but visible aggregates appeared by 10 DAE in various brain regions (Fig. 4H’), demonstrating altered properties of the expanded Htt protein. By 21 DAE, these aggregates became more pronounced and widespread (Fig. 4H”), suggesting progressive accumulation of Htt72Q-Emerald aggregates over time. To test whether Htt72Q-Emerald forms aggregates during development, we examined the brains of third instar larvae expressing Htt72Q-Emerald in neurons from embryogenesis. While Emerald fluorescence confirmed neuronal expression in larval brains, no aggregates were detected (Fig. S4). These findings suggest that expanded Htt expressed during development does not form aggregates at early stages, indicating that the observed neuronal loss at 1 DAE is unlikely due to aggregate formation.

### LactC1C2 improves locomotor performance and prolongs the lifespan of HD flies

To investigate whether masking PS can also protect HD flies from neurodegeneration, we induced LactC1C2-GFP expression alongside the human expanded Htt128Q using the *nsybGal4* driver and monitored the climbing ability and survival rate of HD flies. Remarkably, locomotor performance and lifespan of HD flies expressing LactC1C2-GFP were significantly improved compared to HD only flies, with performance nearly restored to levels observed in control flies (Fig. 4A, B). These results emphasize the critical potential of masking PS exposure to prevent the progression of neurodegeneration.

### LactC1C2 rescues DA and GABA neurons lost only in the adult brain of HD flies

We hypothesized that masking PS by LactC1C2-GFP in the adult brain of HD flies prevented the precocious elimination of stressed neurons by phagocytosis, extending neuronal function. To further investigate the effect of LactC1C2-GFP on neuronal loss, we quantified DA and GABA neurons in the adult brains of HD flies expressing LactC1C2-GFP. At 1 DAE, neuronal loss occurred during development was not rescued by LactC1C2-GFP expression, and the numbers of DA and GABA neurons remained similar between HD model flies and HD flies expressing LactC1C2-GFP (Fig. 4C, D). However, at 21 DAE, a significant rescue of both neuronal populations was observed, with higher counts in HD flies expressing LactC1C2-GFP compared to HD flies, although still lower than in controls (Fig. 4C, D). Notably, the number of DA and GABA neurons at 21 DAE in HD flies expressing LactC1C2-GFP was similar to their numbers at 1 DAE, suggesting that neurons lost during adulthood were rescued, whereas neurons eliminated during development could not be restored (Fig. 4C, D). This difference could be linked to the mode of cell death: cell-autonomous (apoptosis) during development versus non-cell-autonomous (phagoptosis by glia) in the adult. To investigate the basis of this stage specificity, we assessed apoptosis during development (Fig. S5). HD embryos showed increased Dcp-1-positive signal compared to controls, whereas no difference was observed in larval brains, indicating that elevated apoptosis is restricted to embryogenesis. Based on these data, we conclude that in the HD model, PS masking rescues neuronal loss in adulthood but not during development, a difference that we attribute to distinct modes of cell elimination: apoptosis during development and phagoptosis in adulthood.

### Binding to PS is essential for the neuroprotective effect of LactC1C2

To determine whether the neuroprotective effect of LactC1C2 relies on its ability to bind PS, we utilized a mutant version of the protein, LactC1C2-GFP, which lacks PS-binding activity (Fig. 5A, (30). This construct was co-expressed with *skpA* RNAi or Htt128Q in neurons using the *nsybGal4* driver. As shown in Figure 5D and E, flies expressing the mutant LactC1C2-GFP exhibited no significant improvement in motor performance or lifespan compared to *skpA* RNAi-induced or HD model flies (Fig. 5H, I). Furthermore, the numbers of DA and GABA neurons remained similar at 1 DAE and 21 DAE to those observed in the *skpA* RNAi (Fig. 5F, G) or HD model respectively (Fig. 5J, K). These findings indicate that PS binding is essential for the neuroprotective function of LactC1C2-GFP. The data support a model (Fig. 5L) in which LactC1C2 masks PS on the surface of stressed neurons, thereby preventing their recognition and elimination by phagocytic glial cells and promoting neuronal survival (Fig. 5L).

**Figure 5.**
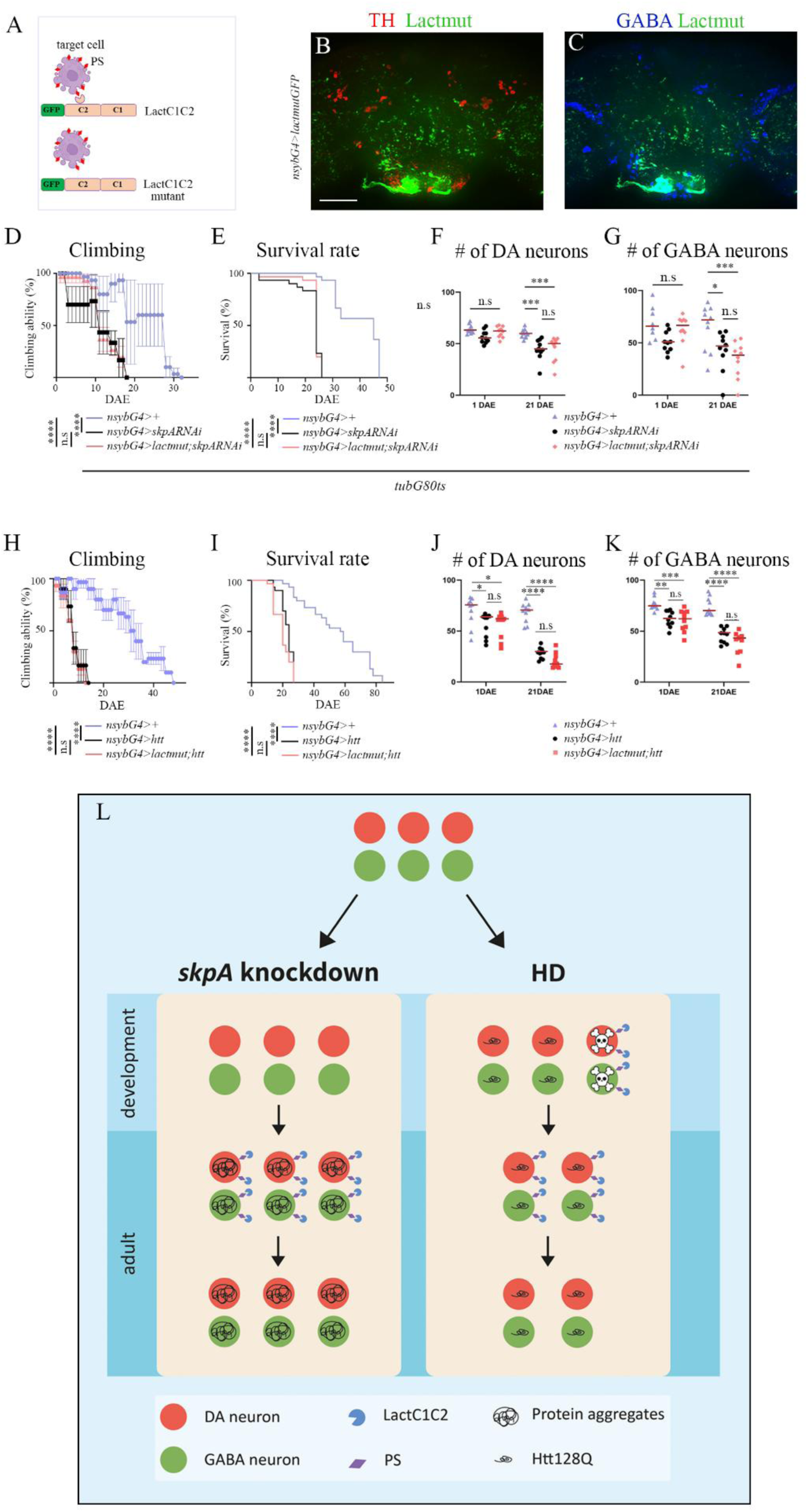
PS binding is essential for the neuroprotective effect of LactC1C2-GFP presented in the proposed model. (A) Schematic representation of LactC1C2-GFP and the mutant form of LactC1C2-GFP (LactC1C2 mutant) unable to bind PS. (B, C) Projections of Z-stacks of the posterior part (∼45 µm) of whole-mount female brain carrying mutant LactC1C2-GFP (*nsybG4>lactmutGFP)* maintained at 29℃ 21 DAE. DA neurons labeled with anti-TH (red), GABA neurons labeled with anti-GABA (blue), mutant LactC1C2-GFP (green). Note homogeneous GFP distribution throughout the brain. (D, E) Climbing ability (D) and survival rates (E) of flies expressing *skpA* RNAi (*tubGal80ts/+; nsybGal4*/*UASskpARNAi*) (black), LactC1C2 mutant and *skpA* RNAi (*tubGal80ts/UASlactmut-GFP;nsybGal4/UASskpARNAi*) (pink) and control flies (*tubGal80ts/+;nsybGal4/+*) (purple). Data are represented as mean ± SEM, n (number of vials, starting with 10 female flies in each) = 3. ****p < 0.0001, n.s = non-significant. (F, G) Mean total number of DA (F) and GABA neurons (G) in dissected brains of flies at 1 DAE and 21 DAE ± SEM, n (number of brains) = ≥8. ***p < 0.001, *p < 0.05, n.s. = non-significant. (H, I) Climbing ability (H) and survival rates (I) of flies expressing human Htt128Q (*nsybGal4/UAShtt*) (black), human Htt128Q and mutant LactC1C2 (*UASlactmut-GFP/+;nsybGal4/UAShtt*) (pink), and control flies (*nsybGal4/+*) (purple). Data are represented as mean ± SEM, n (number of vials, starting with 10 female flies in each) = 3. ****p < 0.0001, n.s = non-significant. (J, K) Mean total number of DA neurons (J) and GABA neurons (K) in dissected brains of flies at 1 DAE and 21 DAE ± SEM, n (number of brains) = 10. ****p < 0.0001, ***p < 0.001, **p < 0.01, *p < 0.05, n.s = non-significant. (L) A proposed model of PS masking as a neuroprotective strategy in two neurodegeneration models. DA and GABA neurons, analyzed in both models, are illustrated as red and green circles, respectively. *skpA* knockdown model: Following normal development, pan-neural adult-specific knockdown of *skpA* results in impaired proteostasis and accumulation of protein aggregates, leading to PS exposure on stressed but viable neurons. Masking PS with LactC1C2 prevents their removal by glial phagocytosis and preserves neuronal integrity. However, neuronal function is not restored, likely due to persistent proteostatic imbalance, which may be treatable within the provided time window. HD model: Pan-neuronal expression of human expanded Htt128Q from embryogenesis leads to early and progressive neuronal loss. In the adult brain, PS masking rescues viable stressed neurons, which retain their function. Neurons lost during development, however, cannot be saved by PS masking with LactC1C2.

### Draper-dependent glial phagocytosis plays model-specific roles in neurodegeneration

We have shown that PS masking is a powerful strategy for preventing neuronal loss in two neurodegenerative models. These findings suggest that blocking phagocytosis by masking the ‘eat me’ signal is sufficient to prevent neurodegeneration. We next investigated whether inhibiting glial phagocytosis through a mechanism distinct from PS masking, by targeting the major phagocytic receptor Draper, could also rescue DA and GABA neurons in both models (Fig. S6). Our results demonstrate that reducing glial phagocytic receptor levels does not fully replicate the protective effects observed using the PS masking strategy. Instead, they indicate that the consequences of impaired glial phagocytosis are highly context-dependent: whereas reduced Draper-mediated phagocytosis is beneficial in the *skpA* RNAi model, it is detrimental in the HD model, likely reflecting distinct roles of glial clearance mechanisms under different neurodegenerative conditions.

## Discussion

In this study, we employed two newly developed *Drosophila* models of neurodegeneration to investigate how glial phagocytosis affects neuronal loss in the diseased adult brain, with a particular focus on the role of PS exposure in this process. In both models, PS masking prevented neuronal loss suggesting that it occurred through phagoptosis where stressed neurons were phagocytosed while still alive. We found that restored functionality of saved neurons depends on the severity and timing of neuronal damage under different neurodegeneration conditions, resulting in the final outcome of their protection. We also reveal the therapeutic potential of truncated Lact as a universal agent to inhibit removal of stressed viable neurons in distinct neurodegeneration contexts. Masking PS has been shown to interfere with the removal of stressed neuronal cells in several cell culture models (29, 39–41). Moreover, an increased number of PS-exposing neurons has been observed in the brains of Alzheimer’s disease patients (42, 43). However, *in vivo* studies examining how PS masking impacts neuronal loss under neurodegenerative conditions remain limited. Here we show that in our first model, adult stage-specific RNAi-induced knockdown of the neuroprotective *skpA,* PS masking with LactC1C2 saved both DA and GABA neurons, suggesting that these neurons were removed via PS-dependent phagoptosis (Fig. 5L). We observed that expression of LactC1C2 partially rescued *skpA* knockdown-induced motor deficits and prolonged lifespan. This implies that although the neurons remained viable, their function may have been compromised and not fully restored. One possibility is that the accumulation of protein aggregates, due to reduced SkpA-mediated proteostasis (19), impaired neuronal function. In addition, other neuronal populations might have been affected by *skpA* knockdown and not rescued by PS masking. Importantly, preventing the clearance of stressed viable neurons provides the time window for potential recovery of neuronal function, an opportunity that would be lost if these neurons were already phagocytosed and degraded.

Our second model, based on pan-neuronal expression of human expanded Htt protein from embryogenesis, revealed that PS masking rescued neuronal loss only in the adult brain of HD flies, but not the loss of neurons during development, maintaining the neuronal counts observed at 1 DAE. These findings suggest that a subset of neurons that had died during development could not be rescued by blocking phagocytic removal. This developmental neuronal loss, triggered by mutant Htt expression, likely occurred through cell-autonomous apoptosis rather than phagoptosis (Fig. 5L). As neuronal apoptosis plays a critical role during CNS development, the apoptotic machinery is readily available in developing neurons and may be activated by mutant Htt. In contrast, inducing neuronal apoptosis in the adult *Drosophila* brain is extremely difficult, as illustrated by the failure of pan-neuronal expression of the pro-apoptotic gene *head involution defective (hid)* to induce detectable cell death at adult stages (Fig. S7). This suggests that phagoptosis is the predominant mechanism of neuronal elimination at this stage. This could explain why PS masking rescues neuronal loss in adults but not during development. Alternatively, mutant Htt may affect developing and mature neurons through distinct mechanisms. Importantly, our findings show that the rescued DA and GABA neurons at the adult stage significantly improved motor performance and extended the lifespan of HD flies to near normal levels, demonstrating that preserved neurons remain functionally competent when spared from glial phagocytosis. Pathogenesis of HD is debated to result from a toxic gain of function of expanded Htt or a loss of wild type Htt function, or both (25). In our HD model, flies still express wild type fly endogenous Htt which may explain the functionality of saved neurons. Interestingly, developmental neuronal death only slightly affected climbing and lifespan of HD flies suggesting that compensatory mechanisms may evolve during development or other functions may be affected, such as memory and learning, which we did not examine in this study. Previous studies in the *Drosophila* olfactory system implicated glial phagocytosis in HD progression via prion-like transmission of mutant Htt aggregates from one cell to another (44–47). Our results complement and expand these findings by revealing a contribution of PS exposure and glial phagocytosis to HD pathogenesis.

We observed that the protective effect of PS masking was not completely recapitulated by silencing the major phagocytic receptor Draper. Reducing glial phagocytic capacity via *draper* knockdown revealed distinct, context-dependent roles for phagocytosis in the adult brain. In the *skpA* RNAi model, decreased Draper levels rescued DA neurons but not GABA neurons, suggesting differential vulnerability of these neuronal populations to Draper levels. The partially improved locomotor performance and extended lifespan are consistent with phagoptosis contributing to the elimination of stressed but viable neurons. In contrast, in the HD flies, reduced Draper levels exacerbated locomotor performance compared to HD flies alone. These findings suggest that Draper-dependent phagocytosis may play an additional protective role in the HD model, potentially by facilitating the clearance of apoptotic neurons generated during pupal stage as well as toxic Htt aggregates or other deleterious material. Together, these results highlight that glial phagocytosis can have opposing functional consequences depending on disease context. Importantly, they suggest that selectively blocking recognition of ‘eat-me’ signals, such as through PS masking, may represent a more targeted strategy to preserve vulnerable neurons while maintaining the clearance of harmful debris.

Our findings in two distinct models of neurodegeneration highlight the therapeutic potential of modulating glial phagocytosis under various pathological conditions. Importantly, masking PS may serve as a universal approach to block glial phagocytosis, since many phagocytic receptors in mammals and flies recognize PS (48). Since in this study we focused on two specific neuronal populations, DA and GABA, our results cannot address whether their loss is responsible for the motor defects and shortened life span or that this phenotype reflects a much larger scale of neurodegeneration that is responsible for the defects.

We found difference in restored functionality of saved neurons across models, suggesting that the severity and/or timing of neuronal damage under distinct neurodegenerative conditions may contribute to the variable outcomes of neuronal protection. We propose that the selective vulnerability of specific neuronal populations in different neurodegenerative diseases depends not only on intrinsic neuronal damage, but in part, on glial phagocytic activity, which may either exacerbate or mitigate disease progression. Preventing PS-dependent clearance of already dead neurons may aggravate disease outcome due to immune response to the uncleared cells; however, rescuing stressed viable neurons may help preserve their function or offer a critical time window for therapeutic intervention as observed here. Our study reveals the truncated form of MFG-E8/Lact as a promising PS-masking agent capable of reducing glial phagocytosis, laying the groundwork for the development of novel therapeutic compounds.

## Materials and Methods

Detailed descriptions of experimental procedures including fly strains, generation of transgenic flies, immunostaining, imaging and quantitative analyses, molecular and biochemical assays, behavioral and survival analyses, and statistical methods are provided in the SI Appendix, SI Materials and Methods.

## Data and code availability

All data generated or analyzed during this study are included in the main text or the supplementary materials. This study did not generate any unique datasets or code.

## Acknowledgments

We are grateful to C. Han, B. McCabe, E. Arama, M. Freeman, O. Schuldiner, the Bloomington Stock Center and the Developmental Studies Hybridoma Bank for fly strains and antibodies. We are grateful to M. Freeman for providing the pUASt-attB-*draperRNAi* plasmid and to M. Volin for initiating the Q-system project. We thank B. Shklyar and L. Simchi from the Bioimaging Unit at the University of Haifa and H. Toledano and B. Shklyar for comments on the manuscript. We also thank the members of the Kurant laboratory for constructive discussions. We gratefully acknowledge financial support from the Israel Science Foundation (grant 274/21).

## Author Contributions

N. K., M. S., E. G., T. F., S. V.-G., M. A and K. H.-M. conducted the experiments; N. K., M. S., E. G. and E.K. designed and N. K., E. G., M. S., T. F., S. V.-G., M. A and K. H.-M. analyzed the experiments; E.K. wrote the paper.

