## Supplemental material for "Masking phosphatidylserine prevents neuronal loss in two distinct *Drosophila* models of neurodegeneration"

#### Supplementary Materials and Methods

##### Fly strains

The following fly strains were used in this work: *w1118* (wild type), *UASlactC1C2-GFP* and *UASlactmut-GFP* (Chun Han), *UASskpRNAi* (#32991 Bloomington Stock Center), *nsybGal4* and *repoQF/TM6B* (Brian McCabe), *UAScytGFP* (##1521 Bloomington Stock Center), *tubQS* (#30130 Bloomington Stock Center), *tubGal80$^{ts}$; TM2/TM6B* (#7019 Bloomington Stock Center), *elavGa4* (Oren Schuldiner), *UAShtt128Q* (#33808 Bloomington Stock Center), *UAShtt25Q-CFP* (#58360 Bloomington Stock Center), *UAShtt72Q-Emerald/TM3* (#58361 Bloomington Stock Center), *UASCD8-GFP* and *UAShid* (Eli Arama), *draper* (M. Freeman), *QUASdraperRNAi* (this study).

For *skpA* RNAi experiments, crosses were maintained at 20°C, and adult flies were transferred to 29°C immediately after eclosion. For HD experiments, both crosses and adult flies were maintained at 25°C. All experiments were conducted using female flies to avoid potential effects of sexual dimorphism.

Quinic acid-containing food was prepared as previously described (49). Briefly, a 1.56 M quinic acid stock solution was prepared by dissolving quinic acid powder (Sigma-Aldrich, catalog no. 77-95-2) in water. For each vial, 100 µl of the quinic acid stock solution was evenly distributed into 10 holes made in 1 ml of food. The food was dried overnight at room temperature and stored at 4 °C for up to two weeks. Newly eclosed female flies were transferred to quinic acid-containing food and flipped onto fresh food every other day.

##### Molecular cloning and transgenic flies

The *draper* RNAi fragment was isolated from the **pUAST-attB** plasmid containing *draper* RNAi (50) and inserted into the **XbaI and SacII** restriction sites of the **QUAS-attB plasmid.** Transgenic flies were generated by BestGene.

**Immunohistochemistry**

For immunohistochemistry, whole brains were dissected, fixed, and stained according to standard procedures. Embryos were collected at stage 16 (13-16h old), fixed and stained according to standard procedures. The following antibodies were used (at the dilutions listed): mouse anti-Draper (#8A1-s1, 1:100) and anti-Eve (#2B8-s, 1:100) from Developmental Studies Hybridoma Bank, mouse anti-TH (1:200; Millipore), rabbit anti-GABA (1:250; Sigma), rabbit anti-cleaved caspase 3 (Dcp-1) (1:100 in embryos, 1:200 in brains), Cell Signaling), chicken anti-GFP (1:100, Roche). Fluorescent Alexa 488, Cy3 or 647 secondary antibodies from Jackson ImmunoResearch were used at 1:200 dilution. Then, brains or embryos were placed in mounting medium (80% Glycerol in PBS). For immunohistochemistry and quantification of DA and GABA neurons, apoptotic cell volume and Draper levels, flies and larvae were dissected at selected ages (DAE).

**Annexin V and LysoTracker staining**

Dissected, unfixed brains were incubated in PBS containing fluorescent Annexin V (final concentration 20ng/ml) or 100 nM LysoTracker Red DND-99 for 10 minutes to label PS exposure and acidic compartments, respectively. Following staining, brains were washed three times in PBS (10 minutes each) and then fixed in 4% paraformaldehyde (PFA) in PBS for 20 minutes. After fixation, samples were washed three additional times in PBS (10 minutes each), mounted and imaged using Apotome microscopy.

**Imaging and Quantification Procedure**

Images of stained brains were acquired on a Zeiss Axio Observer microscope equipped with an Apotome system using the AxioVision software and on a Spinning Disc confocal microscope with NIS software. Image analysis was performed using Zeiss ZEN and Imaris (Oxford Instruments) software. To quantitate the number of DA and GABA neurons, stacks were acquired from the posterior part (~45 µm) of adult brains. To assess levels of PS (Annexin V), expression intensity of Draper (anti-Draper) and phagocytic activity (LysoTracker), microscope stacks from entire brains were analyzed.

Imaging of CFP-tagged Htt25Q and Emerald-tagged Htt72Q was conducted using a Spinning Disc confocal microscope and NIS software. Brains from female flies expressing the respective tagged Htt forms in neurons were dissected and fixed prior to imaging.

**Western Blot Analysis**

*Drosophila* female head extracts were prepared in lysis buffer containing 50 mM Tris-HCl, pH 7.5, 150 mM NaCl, 0.5% Triton X-100, supplemented with a mixture of protease inhibitors (Roche), using a hand homogenizer. Protein concentrations were determined (Bradford reagent, Bio-Rad), and 100 mg proteins were separated by SDS-PAGE (200 V). After electrophoresis, proteins were transferred to a Nitrocellulose Blotting Membrane 0.45 µm (100 mA, 15 V) (Bio-Rad) using a semi-dry blot apparatus. Membranes were blocked using blocking buffer and incubated overnight at 4°C with the rabbit anti-SkpA (1:1000) (Genscript) or mouse anti-Actin

(1:1000) (MP Biomedicals). Membranes were incubated with horseradish peroxide-conjugated secondary antibodies (1:10000) (Jackson ImmunoResearch). Antibody binding was revealed using WESTAR ANTARES (Cyanagen). A LAS4000 luminescent image analyzer (Fujifilm) was used for visualization and image acquisition. ImageJ program was used for image analysis.

##### Climbing ability assay

The climbing assay reflects the locomotor ability of flies during adulthood. We followed climbing activity of 30 adult flies, 10 flies per vial, on a daily basis, by counting the number of flies climbing a distance of 7 cm from the bottom to the top of the vial, within 10 seconds. The flies' medium was replaced every two days until all examined flies lost locomotive skills. At least three independent experiments were performed with independently derived transgenic flies for each genotype.

##### Survival rate assay

The survival rate of flies reflects their lifespan. We measured the survival rate of flies by counting the number of living flies out of 10 flies in each vial, every day after eclosion until the last fly died. In each biological replicate 30 female flies were analyzed, 10 flies per vial, three vials. To avoid sex-related differences we examined only female flies, which were collected as virgins and separated from males. Regular molasses-based food was used on which flies were flipped every other day except of using quinic acid-supplemented food for removing QS suppression.

**Statistical Analysis**

Each experiment was repeated independently a minimum of three times, error bars represent the standard error of replicate experiments. DA and GABA neurons were counted in the designated area of ≥ 10 dissected brains of each genotype. Levels of Annexin V, Draper and LysoTracker volume were measured using Imaris software.

Statistical significance of cell numbers, Dcp-1 and LysoTracker volume data was calculated with Two-way ANOVA followed by Tukey's multiple comparisons test. Draper staining intensity and levels of Annexin V were calculated with Two-way ANOVA followed by Sidak's multiple comparisons test. P values of $< 0.05$ = *, $< 0.01$ = **, $< 0.001$ = ***, $< 0.0001$ = **** were considered significant. P values are indicated in figure legends.

Statistical significance of climbing assay was analyzed employing Two-way ANOVA following a Tukey's post-hoc and Wilcoxon rank-sum test (Mann–Whitney U test). Survival rate was analyzed employing Log rank (Mantel-Cox test) using GraphPad Prism version 8.4.3 for Windows, GraphPad Software, SanDiego, California, USA, https://www.graphpad.com.

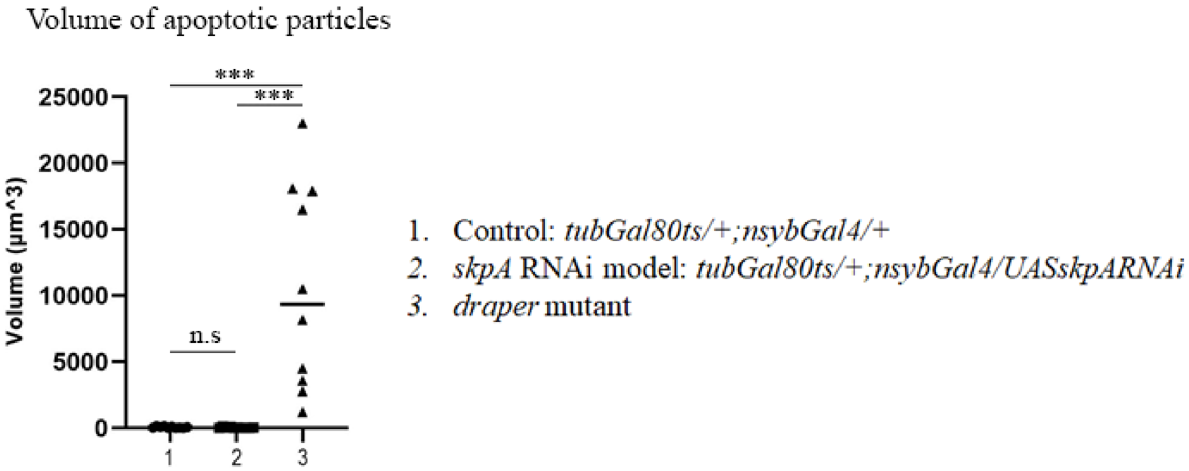


**Figure S1. No detectable apoptosis in *skpA* RNAi knockdown brains.** Mean total volume of apoptotic particles labeled with anti-Dcp-1 antibodies within the central brain area of 14 DAE female flies ± SEM, n (number of brains) = 10. Asterisks indicate statistical significance versus control, as determined by Two Way ANOVA, ***p<0.001, n.s = non-significant.

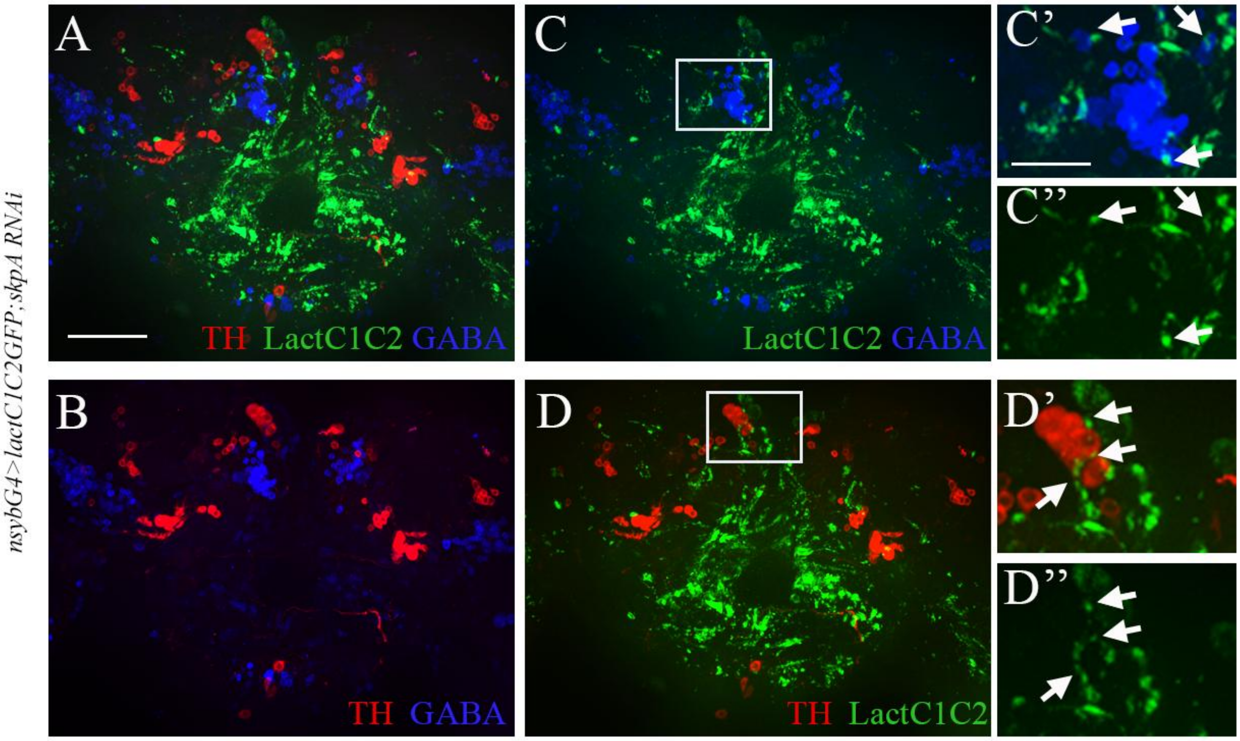


**Figure S2. LactC1C2-GFP is detected on the surface of DA and GABA neurons in the *skpA* RNAi model.** (A - D) Maximum intensity projections of confocal Z-stacks from the posterior region (~45 µm) of whole-mount female brains from the *skpA* RNAi model flies expressing LactC1C2-GFP (*tubGal80ts/UASlactC1C2-GFP;nsybGal4/UASskpARNAi*) maintained at 29 °C for 10 DAE. DA neurons are labeled with anti-TH (red), GABA neurons with anti-GABA (blue) and LactC1C2-GFP is shown in green. (C', C'', D', D'') Magnified views of boxed regions in (C) and (D) shown as separate channels. Arrows indicate TH- and GABA-labeled neurons with GFP signal accumulating on their surface. Bar, 20 µm.

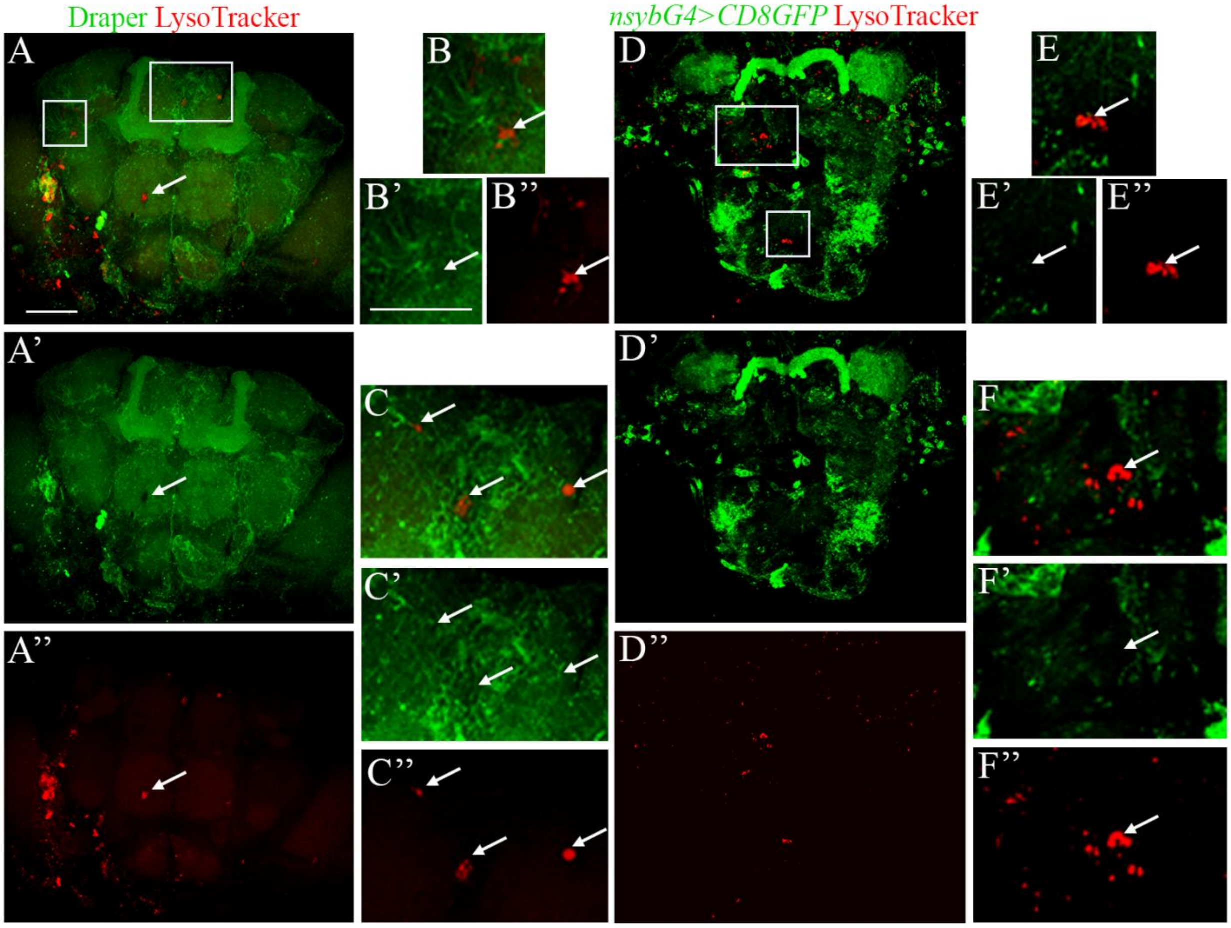


**Figure S3. In *skpA* knockdown brains LysoTracker signal is detected in glial cells.** Representative projections of Z-stacks of whole-mount brains from female flies maintained at 29°C for 16 days (16 DAE). LysoTracker (red). (A-C") Glial membranes labeled with anti-Draper (green). (D-F") Neuronal membranes labeled with *nsybGal4>CD8GFP* (green). Bar, 50 µm. Two rectangles in (A) and (D) are enlarged and shown in (B-C") and (E-F") respectively. Arrows indicate Lysotracker signal surrounded by glial membranes (B-C") but not by neuronal membranes (E-F").

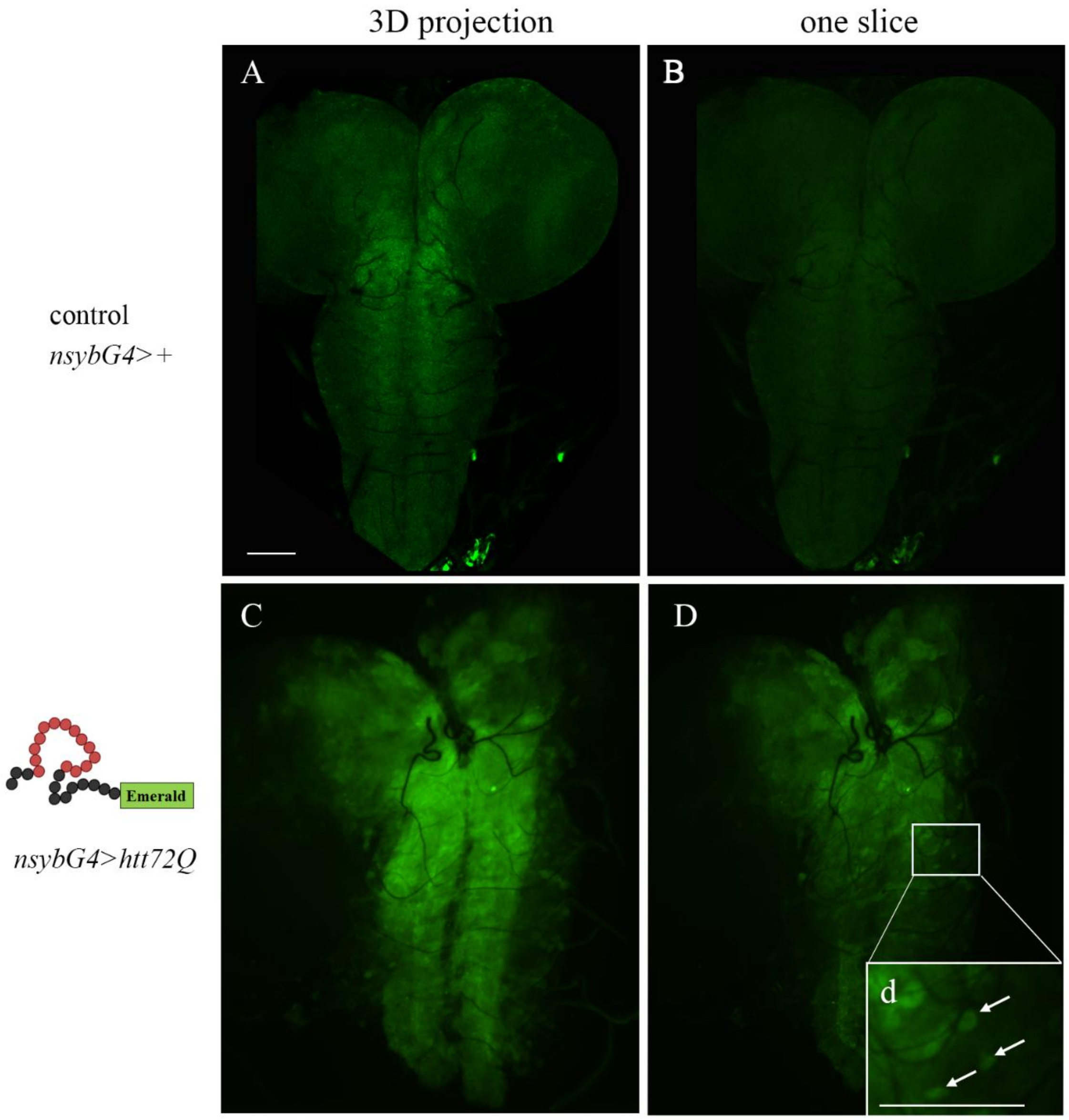


Figure S4

**Figure S4. Htt72Q-Emerald does not form aggregates in the larval brain.** (A, C) Maximum intensity projections of confocal Z-stacks and (B, D) single optical sections from whole-mount third instar larval brains. (A, B) Control (*nsybGal4/+*). (C, D) HD model expressing Htt72Q-Emerald (*nsybGal4/UAShtt72Q-Emerald*). (d) Magnified view of the boxed region in (D). Arrows indicate Emerald fluorescence without detectable aggregate formation. Bar, 100 µm.

**DA and GABA neurons die through apoptosis during development of HD flies.**

To examine whether DA and GABA neurons, which were not rescued by PS masking, were eliminated through apoptosis during development, we performed anti-Dcp-1 staining and quantified apoptotic cell volume using Imaris software in embryos and third instar larval brains of HD model and control. A larger volume of Dcp-1-positive apoptotic particles was found in HD model embryos compared to controls (Fig. S5A-A”). In contrast, no difference in apoptotic cell volume was observed between HD and control larval brains (Fig. S5D-D”). These results suggest that excess apoptosis induced by neuronal expression of Htt128Q occurs primarily during embryogenesis rather than at larval stages.

To further characterize neuronal apoptosis and determine whether DA and GABA neurons die during embryogenesis, we stained embryos with anti-TH and anti-GABA antibodies. While anti-TH staining was successful (Fig. S5B-B”), anti-GABA antibodies, which work well in larval and adult brains, failed to label embryonic neurons. Therefore, we quantified an additional neuronal population marked by anti-Eve antibodies and found no difference between control and HD embryos (Fig. 5SC-C”), suggesting that DA and Eve-positive neurons do not undergo apoptosis during embryogenesis in the HD model. Consequently, no conclusion can be drawn regarding GABA neurons at this stage.

However, when counting DA and GABA neurons in larval brains, we observed a reduced number of GABA neurons in HD model larvae compared to controls (Fig. 5SF-F”), suggesting that GABA neurons likely die through apoptosis during embryogenesis, consistent with the increased Dcp-1-positive volume observed at this stage (Fig. 5SA-A”). In contrast, DA neuron numbers were unchanged in both HD embryos and larvae (Fig. 5SB”, E”), indicating that DA neurons do not die during embryonic or larval stages. Since we detect a reduced number of DA neurons in emerging adults, we propose that DA neuronal loss in the HD model likely occurs during pupal stages. Altogether, these results demonstrate selective and stage-specific vulnerability of DA and GABA neurons to Htt128Q expression in the HD model.

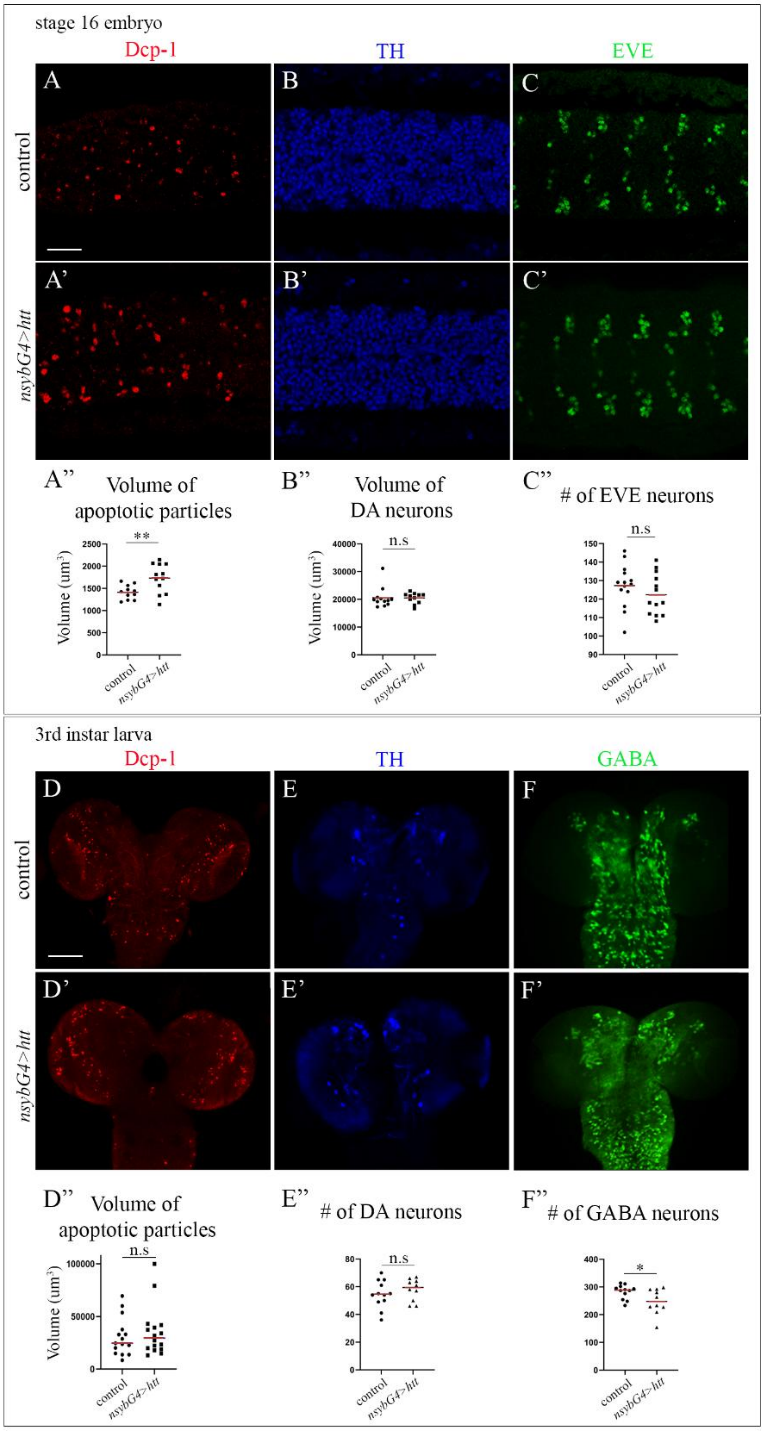

**Figure S5. DA and GABA neurons die through apoptosis during development of HD flies.** (A-C') Imaging of embryos was conducted using Zeiss LSM 980 confocal microscope. Microscope Z-stacks were acquired from the neural cortex of the embryonic ventral nerve cord. Projections of confocal stacks from stage 16 embryonic CNS (ventral view). (A, A') Apoptotic cells labeled with anti-Dcp1 (red). (B, B') DA neurons labeled with anti-TH (blue). (C, C') Eve-positive neurons labeled with anti-Eve (green). (A-C) Control (*nsybGal4/+*). (A'-C') HD model (*nsybGal4/UAShtt*). Bar, 20 µm. (A") Mean total volume of apoptotic particles. (B") Mean total volume of DA neurons. (C") Mean total number of Eve-positive neurons within CNS sections (22 sections; total thickness 7.7 µm; 5 segments) ± SEM; n (number of embryos) > 11. Asterisks indicate statistical significance versus control as determined by Student's t-test, **p < 0.01, n.s = not significant. (D-F') Projections of confocal stacks from third instar larval brains. (D, D') Apoptotic cells labeled with anti-Dcp1 (red). (E, E') DA neurons labeled with anti-TH (blue). (F, F') GABAergic neurons labeled with anti-GABA (green). (D-F) Control (*nsybGal4/+*). (D'-F') HD model (*nsybGal4/UAShtt*). Bar, 20 µm. (D") Mean total volume of apoptotic particles. (E") Mean total volume of DA neurons. (F") Mean total number of GABA neurons within one entire lobe of third-instar larval brain ± SEM; n (number of samples) ≥ 10. Asterisks indicate statistical significance versus control as determined by Student's t-test, *p < 0.05, n.s = not significant.

**Draper-dependent glial phagocytosis plays model-specific roles in neurodegeneration**

To investigate whether inhibiting glial phagocytosis through a mechanism distinct from PS masking, by targeting the major phagocytic receptor Draper, could also rescue DA and GABA neurons in both models (Fig. S6), we analyzed neurodegeneration in both the *skpA* RNAi and HD models when phagocytosis was reduced by decreasing Draper expression. Because *draper* null mutant flies exhibit a severe neurodegenerative phenotype due to the accumulation of apoptotic debris in the brain during development (39), Fig. S6D'), we employed an inducible RNAi-mediated knockdown of *draper* in adult glia (Fig. S6A). Specifically, we generated transgenic flies carrying an inducible QS/QF system, in which QS-mediated repression can be relieved by the addition of quinic acid to the food (40, 41), Fig. S6A). To validate this approach, brains were stained with anti-Draper antibodies to confirm Draper reduction (Fig. S6B, D-E"') and with anti-Dcp-1 antibodies to detect apoptotic particles (Fig. S6C-E"'). Consistent with previous observations in *draper* null mutants (Fig. S6D'), continuous *draper* knockdown throughout development using *repoQF* resulted in the accumulation of numerous apoptotic particles in the brain (Fig. S6D). In contrast, very few apoptotic particles were detected in the absence of *draper* RNAi induction (Fig. S6C) or when QS-mediated repression remained active throughout development (Fig. S6D"). Importantly, brains in which QS repression was relieved only after eclosion by transferring newly emerged adults to quinic acid-supplemented food exhibited almost no apoptotic particles (Fig. S6D"'). These findings confirm that the accumulation of apoptotic corpses results from impaired Draper function during development.

**Glial knockdown of *draper* rescues DA but not GABA neurons and improves locomotor performance and lifespan in *skpA* RNAi flies**

We first assessed the effect of reduced Draper in the *skpA* RNAi model by analyzing flies in which *draper* was knocked down specifically in adulthood in combination with neuronal *skpA* knockdown (Fig. S6F-I). Locomotor performance and lifespan were improved in *skpA* RNAi flies with adult-specific *draper* knockdown compared to the *skpA* RNAi alone (Fig. S6F, G). Notably, reducing Draper levels in adult glia prevented the loss of DA neurons (Fig. S6H) but not GABA neurons (Fig. S6I), suggesting differential vulnerability of these neuronal populations to reduced Draper-mediated phagocytic activity. These results indicate that reducing glial Draper in the adult neurodegenerating brain can also ameliorate functional decline and extend survival in this model. We however notice that PS masking by LactC1C2 provides better protection by rescue not only DA but also GABA neurons.

**Glial knockdown of *draper* aggravates motor impairment without affecting lifespan in HD flies**

We then examined the effect of glial adult stage-specific knockdown of *draper* in the HD flies (Fig. S6J-M). Locomotor performance was significantly worst in these flies compared to the HD alone (Fig. S6J). We observed that the number of DA and GABA neurons at 1 DAE was similar to the HD flies alone, confirming that reducing Draper levels only in adult glia does not affect neuronal loss during development (Fig. S6L, M). Importantly, adult stage-specific *draper* knockdown did not significantly affect lifespan (Fig. S6K) but increased neuronal number at 21 DAE (Fig. S6L, M).

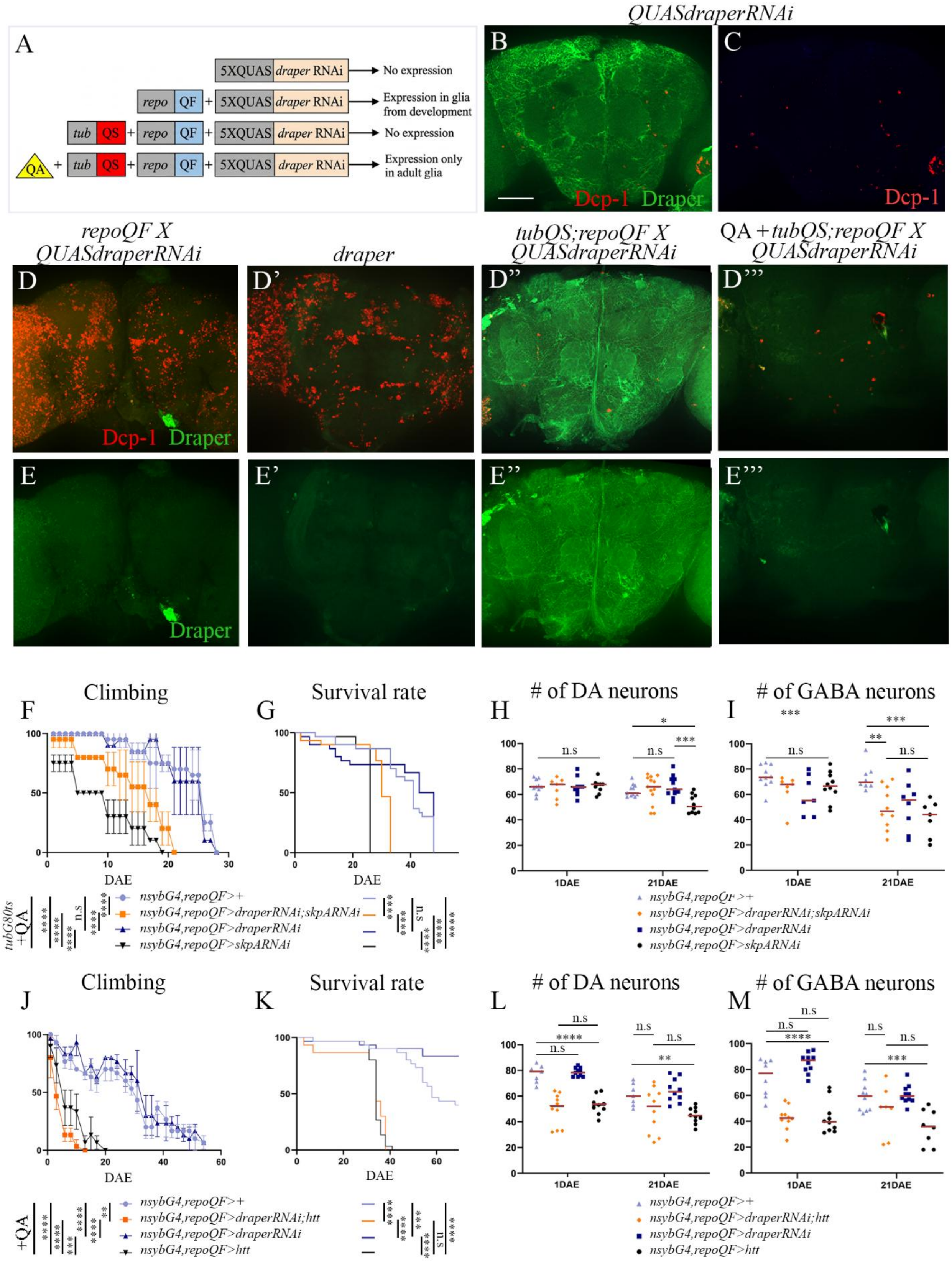
QUASdraperRNAi
A
5XQUAS draper RNAi
No expression
repo QF + 5XQUAS draper RNAi
Expression in glia from development
tub QS + repo QF + 5XQUAS draper RNAi
No expression
QA + tub QS + repo QF + 5XQUAS draper RNAi
Expression only in adult glia
B
Dcp-1 Draper
C
Dcp-1
repoQF X QUASdraperRNAi
draper
tubQS;repoQF X QUASdraperRNAi
QA + tubQS;repoQF X QUASdraperRNAi
D
D'
D''
D'''
Dcp-1 Draper
E
E'
E''
E'''
Draper
F
Climbing
G
Survival rate
H
### of DA neurons
I
### of GABA neurons
DAE
1DAE
21DAE
n.s
tubG80ts +QA
nsybG4,repoQF>+
nsybG4,repoQF>draperRNAi;skpARNAi
nsybG4,repoQF>draperRNAi
nsybG4,repoQF>skpARNAi
J
Climbing
K
Survival rate
L
### of DA neurons
M
### of GABA neurons
+QA
nsybG4,repoQF>+
nsybG4,repoQF>draperRNAi;htt
nsybG4,repoQF>draperRNAi
nsybG4,repoQF>htt

**Figure S6. Reduced Draper differentially affects neurodegeneration in the *skpA* RNAi and HD models.** (A) Schematic representation of transgenic constructs used for Q-system–induced glia-specific *draper* RNAi knockdown. (B-E''') Confocal images of the posterior region (~45 µm) of whole-mount female brains immunostained with anti-Draper (green) and anti-Dcp-1 (red). (B, C) Control: *QUASdraperRNAi* without QF. (D, E) *QUASdraperRNAi/+;repoQF/+* (expression initiated during embryogenesis). (D', E') *draper* mutant. (D", E") *tubQS/+;QUASdraperRNAi/+;repoQF/+* (no quinic acid). (D''', E''') *tubQS/+;QUASdraperRNAi/+;repoQF/+* (quinic acid-treated food). (F, G, J, K) Climbing ability (F, J) and survival rates (G, K) of control flies (purple), *skpA* RNAi model flies (F, G) or HD model flies (J, K) (black), model + *draper* RNAi (orange), and *draper* RNAi only (blue). Data are shown as mean ± SEM; n (number of vials, each initially containing 10 female flies) = 3. HD survival experiments were performed for 70 days and were terminated before all flies in the control and *draper* RNAi-alone groups had died. ****p < 0.0001, ***p < 0.001, **p < 0.01, n.s = non-significant. (H, I, L, M) Mean total number of DA neurons (H, L) and GABA neurons (I, M) in dissected brains at 1 and 21 DAE ± SEM; n (number of brains) = ≥7. ****p < 0.0001, ***p < 0.001, **p < 0.01, n.s = non-significant.

***skpA* RNAi-induced model genotypes:**

Control: *QS/+;tubG80ts/+;nsybGal4,repoQF/+*

Model + *draper* RNAi:

*QS/+;tubG80ts/UASQUASdraperRNAi;nsybGal4,repoQF/UASskpARNAi*

*draper* RNAi only: *QS/+;tubG80ts/UASQUASdraperRNAi;nsybGal4,repoQF/+*

Model only: *QS/+;tubG80ts/+;nsybGal4,repoQF/UASskpARNAi*

**HD model genotypes:**

Control: *QS/+;;nsybGal4,repoQF/+*

Model + *draper* RNAi: *QS/+;UASQUASdraperRNAi/+;nsybGal4,repoQF/UAShtt*

*draper* RNAi only: *QS/+;UASQUASdraperRNAi/+;nsybGal4,repoQF/+*

Model only: *QS/+;;nsybGal4,repoQF/UAShtt*

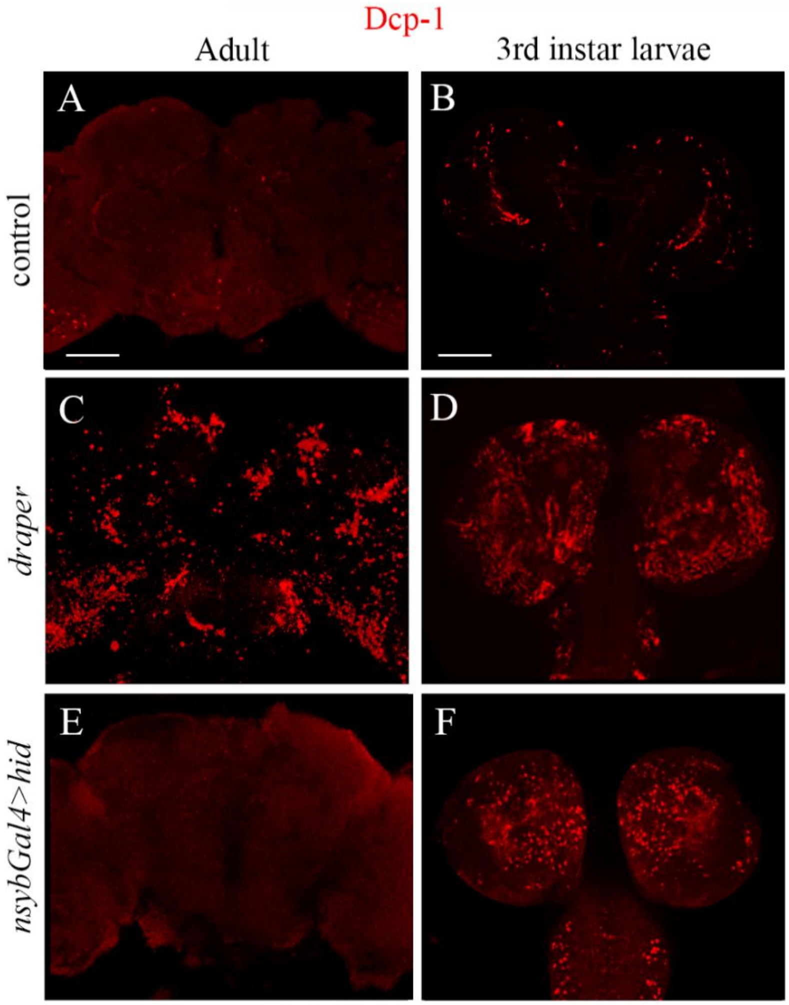


**Figure S7. Neuronal expression of the pro-apoptotic protein Hid induces massive apoptosis in larval brains but not in adult brains.** (A–F) Confocal images of whole-mount (A, C, E) adult brains (7 DAE) and (B, D, F) third instar larval brains stained with anti-Dcp-1 to detect apoptosis. (A, B) Control (*tubGal80ts/+; nsybGal4/+*). (C, D) *draper* mutant. (E, F) *tubGal80ts/UAS-hid; nsybGal4/UAS-hid*. All crosses were maintained at 20°C to prevent *hid* expression during development. (A, C, E) Newly eclosed females were shifted to 29°C, and brains were dissected at 10 DAE. (B, D, F) Third instar larvae were kept at 29°C for 24 hours prior to dissection. Note the absence of detectable apoptotic signal in adult brains expressing *hid* pan-neuronally (E) compared to the strong apoptotic signal in larval brains (F). Bar, 50 µm.
